# Oversharing by honey bees and the spread of viruses

**DOI:** 10.1101/2022.05.15.492017

**Authors:** Zachary S. Lamas, Eugene V. Ryabov, David J. Hawthorne, Jay D. Evans

**Author notes:** **Correspondence:** Zachary S. Lamas.

## Abstract

Deformed wing virus (DWV) is an emerging insect pathogen efficiently transmitted through communicable and vector-borne routes with *Apis mellifera*. Continual transmission of DWV between hosts and vectors is required for maintenance of the pathogen within the population, and this vector-host-pathogen system offers unique disease transmission dynamics for pathogen maintenance between vector and a social host. In a series of experiments, we study how vector-vector, host-host and host-vector transmission routes maintain DWV in a honey bee population. We found co-infestations on shared hosts allowed for movement of DWV from mite to mite. Additionally, two social behaviors of the honeybee, trophallaxis and cannibalization of pupae, provide routes for communicable transmission from bee to bee. Communicable circulation of the virus solely amongst hosts was then shown to act as a reservoir of DWV for naïve *Varroa* to acquire and subsequently vector the pathogen. Our findings illustrate communicable transmission between hosts can maintain and influence the future acquisition and vectoring of a pathogen by a vector. There are a handful of other infectious diseases, both zoonotic and which impact human health, that have potentially similar transmission dynamics.

## Introduction

Continual transmission between infectious and susceptible individuals is essential for the persistence of a pathogen in a population^1^. Infectious individuals, aptly called maintenance hosts, form the host reservoir. Over the last 60 years, numerous theoretical models have been developed to describe the transmission dynamics of infectious diseases^2,3^, ultimately encompassing both vector-host-pathogen systems and the threats posed by emerging diseases^4,5^. These models rely on complete and accurate estimates of routes and parameters of transmission, which may be lacking in emerging diseases. One such disease is caused by the reemerging pathogen Deformed wing virus (DWV). Once rarely observed in its honeybee host *Apis mellifera*, DWV has become highly prevalent in managed and wild colonies of honeybees^6^, and detected in more than 65 arthropod species, presumably an indicator of spillover from infected honeybees^7^. DWV is an RNA virus from the Iflaviridae family that is transmitted both communicably and as a vector-borne pathogen in global honey bee populations^*6*^. This virus was present, but had low prevalence in honeybee populations prior to the introduction of *Varroa destructor*, an ectoparasitic mite and competent vector of honeybee viruses that jumped hosts from *Apis cerana* to *Apis mellifera*^*8*^. Impacts of DWV garnered global attention when severe losses in managed and feral honeybee colonies were reported in 2006 in North America^9^, leading to intensive research on DWV and its impacts^7^.

Honeybees are the primary host of DWV, with other Hymenoptera serving as dead end hosts for this virus^6^. DWV is transmitted via multiple communicable modes among the thousands of members found in honeybee colonies^10^. Mockel *et al* showed that adult workers can develop covert (subclinical) infection after oral exposure to the virus^11^. Later, consumption of infectious pupae by adult bees followed by spread through trophallaxis between nestmates was shown to be an efficient route for transmission, which also resulted in covert infection^12^. Brood rearing is an essential function in a honeybee colony. Yue and Genersch detected DWV in larval food, suggesting brood rearing may be a route of infection for developing bees^13^. In addition, infection in developing brood has been shown to originate through the queen via vertical transmission. Amiri *et al*. demonstrated experimentally 1/3 of colonies had eggs infected with DWV. The virus was deposited on the surface of the eggs indicating transovum transmission^14^. Direct infection to queens has been demonstrated experimentally through artificial insemination of virgin queens and in field studies through natural mating^15,16^. Importantly, covert infections arising from the queen and passing vertically to her thousands of offspring provide for sustained transmission within the colony, a key parameter in the maintenance of DWV. De Miranda and Fries postulated that infections through this route would influence *Varroa*-mediated and fecal-oral-cannibalization transmission routes by potentially infecting a large swath of bees^17^.

*Varroa* as an active vector of honey bee viruses is a recent phenomenon. *Varroa* feed on bees both during their reproductive phase on a developing larva, and during a dispersal phase on adult bees^18-20^. Feeding bouts during both periods offer opportunities for vector-mediated transmission to hosts. Overt (clinical) infections are most evident in the form of crippled wings, shortened abdomens, and adults with motor-paralysis resulting from transmission during larval development^9,21^. Subclinical infections occur when bees are parasitized during development but do not develop physical deformities, and are often associated with lower viral loads^9^,^22^. *Varroa* can be infectious with a virus, or naïve, without a virus^23^. Bee to mite transmission occurs when *Varroa* feed on an already infected host which has developed a high level of infection prior to the *Varroa*’s feeding^20^. Bowen-Walker and Gunn inferred this from the high levels of DWV in *Varroa* collected from infectious bees with crippled wings^24^. Hosts harboring high levels of DWV will be especially infectious for naïve *Varroa* which feed upon them.

Mite-mite transmission has not yet been observed but could potentially occur during both the *Varroa*’s reproductive phase or during the dispersal phase when two or more *Varroa* share the same larval or adult bee host, simultaneously or concurrently. Multiple infestations are not as common as single infestations, but may be an important feature in the maintenance and spread of DWV in the *Varroa* population^25^. Similar features are important in the maintenance of pathogens in other parasites and vectors, most notably with cofeeding ticks^26-28^. Both mite-mite and bee-mite transmission routes could be important in the maintenance of DWV in the *Varroa* population.

In a series of experiments, we asked if transmission occurs mite to mite on shared hosts, and through bee to mite routes. When examining bee to mite transmission routes we asked if communicable transmission of a virus among honeybees can influence acquisition of the virus by mites and future vector transmission of the virus. We show a surprising degree of interaction between communicable, and vector borne transmission modes; social behaviors of the host species aid in transmission, acquisition and future vectoring of the pathogen by the introduced parasite. In the most notable example, hygienic removal of diseased pupae by adult bees, a form of social defense in honeybees, aids in the spread and persistence of DWV in hosts and *Varroa*. This study integrates these routes, showing that honey bees themselves can amplify the impacts of a new vector.

## Materials and Methods

### Construction of custom cages

Custom laboratory arenas were constructed from acrylic sheets so that bees could be physically isolated in adjoining, but divided chambers. The divider was a solid piece of acrylic with a 7/8 inch hole cut in the center. The hole was sealed with 1/8 inch hardware cloth. This served as a trophallaxis port in which bees could orally exchange food with limited physical contact. 1/8 fiberglass mesh screen was used for ventilation at the end of each chamber. The interior dimensions of each cage were 2.5” x 2.5” x 2.75”.

### Viral inoculum used for studies

A cDNA clone derived variant of DWV-A tagged with nanoluciferase (NLuc) gene was used throughout the trials^29^. This virus has two inserted genetic tags in its nucleotide sequence which allow for reliable detection and made it possible to distinguish the clone-derived virus from wild-type variants. The nucleotide sequence of NLuc reporter inserted in the viral genome made it possible to detect and quantify the virus RTqPCR. The presence of second modification, the site for unique rare-cut site for PacI restriction endonuclease that can be verified by digestion of RT-PCR product^29^. In this way, dual verification of the presence of the tagged virus was achieved. For the rest of the document inoculum refers to this as the DWV-A NLuc virus. Stock samples were provided by Evans and Ryabov and maintained at the USDA-ARS Bee Research Laboratory in Beltsville, Maryland.

### Molecular preparation and qPCR

All samples were extracted using standard techniques^30^. Whole bee specimens were extracted in trizol and then RNA was used to produce cDNA using single reaction reverse transcriptase according to manufacturer specifications (BioRad, IScript). Total viral cDNA was quantified using real time qPCR and a 10 fold dilution series of prepared standards exactly as described in Posada-Florez et al.^12^.

Verification of 5′ region of DWV RNA (30–1266 nt) containing the PacI site was performed through digestion of the Pac1 and then visualization on an agarose gel^29^.

### Experiment 1: Can communicable transmission of DWV between honeybees amplify future acquisition and vectoring of the pathogen?

#### Phase 1: Establishing infectious donor bees

White-eyed pupae were injected with 1 ul of either viral inoculum (10^7^ GE) or PBS and incubated for 12 hours at 34 °C and 40% humidity. *Varroa* were sampled from adult bees collected from a moderately infested colony not exhibiting any signs of overt varroosis or brood disease. *Varroa* were serially passaged for five days on pupae procured from a healthy colony that did not yield any *Varroa* detections (5 mites per pupa host). *Varroa* were randomly assigned to a control (PBS injected) or experimental (viral injected) pupae for their last passage. At 12 hours after injection, pupae were transferred into a 00-sized gel cap. *Varroa* (4 to 7) were then placed inside the gel cap with the pupae, and returned to the incubator to feed for 24 hours (36 hours since injection).

After feeding for 24 hours on injected pupae, *Varroa* were transferred to 3 day old adult bees. Adult bees were procured from a frame of emerging brood sourced from a healthy colony exhibiting no clinical signs of disease that yielded no *Varroa* detections via an alcohol wash. Bees and *Varroa* were incubated at 34°C at 40% humidity, and checked twice daily for *Varroa* feeding. The bee was removed from the cage with a pair of soft tweezers when a *Varroa* was observed in a known feeding position. The bee was held upside down by the wings or thorax with one hand, while the researcher agitated the *Varroa* from the feeding position with a Chinese grafting tool (Mann Lake, Hackensack, MN). Once removed, the *Varroa* was scooped from the bee and stored at -80 °C. The bee was thoroughly inspected for additional *Varroa*, and then transferred to a “donor” compartment of the custom arenas. Bees were added in this manner to the arenas over the course of 4 days and incubated for a total of 7 days (from first addition). Bees were checked twice daily for the presence of *Varroa*, and to record bee mortality.

#### Phase 2: Addition and maintenance of recipient bees

At day 8 of the trial, recipient bees were added to the second compartment of the custom arenas. Recipient bees were newly emerged and procured in the same way as the donor bees. Recipient bees were thoroughly inspected for *Varroa* at emergence before introduction into the custom arenas, and inspected twice daily thereafter. Beginning on the second day after introduction of recipient bees trophallaxis was encouraged by removing sucrose feeders from the recipient’s side. Sucrose feeders were removed for four hours, after which bees were checked for *Varroa*, and then feeders were returned. This continued for 8 more days (15 total days since first addition of donor bees). *Varroa* that were not experimentally inserted were not observed in any of the replicates during the trial.

#### Phase 3: Removal of donor bees and addition of *Varroa*

On day 16, the recipient bees were removed from the arenas and stored in at - 80 °C. Recipient bees were removed by lightly chilling the whole arena at 4°C for 10 minutes. The lid of the recipient compartment was removed, and the bees were knocked quickly into a wide lip plastic cup. A fine mesh net was promptly sealed over the bees to prevent escaping. The cages now with only recipient bees were returned to the incubator.

Recipient bees were thoroughly inspected for *Varroa* one final time (no *Varroa* were found in any of the arenas). *Varroa* were then added to each cage of recipient bees. *Varroa* were procured and serially passaged as previously described on non-injected pupae prior to their introduction onto the adult recipient bees. *Varroa* were allowed to feed on adult recipient bees for 4 days after which *Varroa* were removed as previously described and placed onto a white eyed pupae (non-infested, procured from a healthy colony) in 00 gel caps. *Varroa* and pupae were incubated for 6 days and then preserved at -80 °C for later RNA extraction.

### Experiment 2: Do *Varroa* acquire virus when feeding upon adult bees which previously cannibalized pupae?

Pupae were injected with the viral inoculum as previously described or PBS and then incubated as previously described. *Varroa* were collected and introduced to the pupae as previously described. After 24 hours of feeding on pupae, *Varroa* were removed from the pupae and passaged onto a new pupa. *Varroa* were incubated on their new pupae for 5 days, and then frozen in at -80 °C. These pupae served as the experimental or control pupae for the experiment.

Cages of 30 adult bees were made by collecting worker bees from a healthy colony which did not yield detections of *Varroa* in an alcohol wash. Cages were constructed in exactly the same way and with the same materials as in Chapter 1^20^. Cages were randomly assigned to one of three treatment groups: no pupae, pupae with virus, pupae without virus. One pupa was removed from the -80°C freezer for each experimental cage, thawed and immediately fed to the bees. This was repeated on the following day, after which bees were incubated for an additional 9 days (11 days total since first introduction).

A new series of *Varroa* were procured and passaged as previously described, and then introduced into the experimental cages. *Varroa* were allowed to feed upon these adult bees for 4 days, after which *Varroa* were individually removed and introduced into a 00 gel cap with a white eyed pupa (healthy, non-injected). *Varroa* were allowed to feed on pupae for 48 hours, at which point the *Varroa* were removed and stored at -80 °C. Pupae were incubated for 4 more days and then stored at -80 °C.

### Experiment 3 -4: Do *Varroa* transmit virus via shared honeybee hosts? Collection and marking of *Varroa*

*Varroa destructor* were sourced and hand collected from a heavily infested colony not showing overt signs of *Varroa* parasitism. Adult bees were inspected and those with *Varroa* in their feeding positions were stored in a ventilated cage^20^. Bees and *Varroa* were returned to the laboratory and stored in an incubator at 34 °C and 40% humidity.

*Varroa* were marked in these trials with a small amount of paint applied dorsal-distally directly onto the carapace. Paint was applied with a 0000 fine tip paintbrush (Javis: 4/0 nylon, England) with a small downward strike moving posteriorly from the dorsal tip of the *Varroa*. Paint from fine-tipped oil-based permanent markers were used. Paint was procured by ejecting the tip of the marker onto a plastic surface until a pool of paint formed. The tip of the brush was then dabbed against the surface of the puddle. In this way, a small amount of paint could be applied to the surface of a *Varroa*. Colors were used to identify *Varroa* by group. For example, *Varroa* painted blue were part of the viral group, clearly identifying them differently than the *Varroa* from the control group which were painted orange. The researcher could observe, track and recollect multiple *Varroa* co-housed on the same group of bees.

#### Maintenance

*Varroa* were serially passaged on pupae procured from healthy (no observed *Varroa*) colonies as a means of reducing potential viral loads carried by these *Varroa*^31^. *Varroa* were enclosed with pupal hosts in a labeled 0 gel cap and pupae were changed every 2 days, a method shown to ultimately cleanse *Varroa* of DWV-A. *Varroa* were passaged 3 times prior to the start of the trial. Next, experimental-phase *Varroa* were prepared by passaging on pupae that had been injected with either 10ul of PBS or PBS with viral inoculum 12 hours prior to *Varroa* introduction. These *Varroa* were incubated in 0 gel caps with their respective pupa for 24 hours (total 36 hours post injection), prior to their use in the below trials.

### Experiment 3- Examining potential *Varroa* to *Varroa* transmission on shared pupal hosts

A frame of recently capped worker brood was removed from a healthy *Varroa* free colony and brought to the laboratory. The corner of a cell capping was lifted with a razor blade so that *Varroa* could be introduced into the recently capped larval cell. *Varroa* were introduced by picking one up with a Chinese grafting tool, and then tipping the tip of the grafting tool with the *Varroa* into the opening of the torn cell. *Varroa* were coaxed into the cell by prodding them with an additional grafting tool held in the other hand. A second *Varroa* was immediately introduced in the same manner. Cell cappings were resealed by pressing the torn capping back down, and using another cell capping as a “bandage”. This was done by removing an entire cell capping with a pair of tweezers, flipping it over, and firmly pressing the waxed side (silk side up) against the “wound” of the experimental cell. If a cell was damaged too greatly to be re-sealed in this way, then it could be capped with a #6 gel cap.

Two *Varroa* were introduced into each cell. Introductions were made by pairings: control x control, or control x virus-exposed (viral). Experimental cells were numbered in chronological order of insertion, with *Varroa* insertion, and the source pupa of each respective *Varroa*. White paint markers were placed to provide a geographical reference point on the surface of the frame, to locate experimental cells. *Varroa* and pupal hosts were recovered 9 days after insertion. Experimental cell locations were confirmed by references cells and the laboratory log. Once confirmed, cell cappings were removed with a pair of tweezers. The pupae were removed and saved at -80°C. *Varroa* were collected from the cells using a Chinese grafting tool, and then inserted with a pink-eyed pupal host in a labeled 0 gel cap. Pupae and *Varroa* were incubated for 48 hours, at which time *Varroa* were removed and stored at -80°C until RNA extraction. Pupae were returned to the incubator for another 4 days, after which samples were preserved at -80°C.

### Experiment 4 Examining mite to mite transmission on shared adult hosts Bee cages

For experiment 4, cages of 8 individually marked bees were used, and established in the same manner in Material and methods, Chapter 1. Marked *Varroa* were placed into prepared cages of marked 3-day-old adult bees. *Varroa* locations were checked 2 hours after introduction and then every 12 hours thereafter for 10 continuous days. Daily *Varroa* and host mortality was recorded. All samples were saved at -80°C until extraction. The location of each *Varroa* was recorded every 12 hours, recording whether the *Varroa* was feeding or not on the host bee, and specifically which host bee. Recordings were made in a way to differentiate each *Varroa* from the other, and to track them independently.

### Statistical Analysis

All statistical analysis were performed in RStudio using BaseR and associated packages. For all trials, count data were used to report the observed number of infectious bees or **Varroa** over the number of non-infectious bees or **Varroa** in a trial. In order to test viral load differences in exposed and non-exposed Varroa and bees to the NLuc virus an ANOVA was performed to see if there were significant differences between the groups. The residuals of data were visualized and then tested for normality using the Shapiro-Wilk test. DWV-A titers were compared across groups using an ANOVA. When the assumptions of normality were not met with NLuc titers a non-parametric Kruskal-Wallis ANOVA was used. Statistical significance was set at P < 0.05.

## Results

### Experiment 1 Can *Varroa* acquire virus following communicable transmission between honeybees?

Using tagged virus clones, we showed that *Varroa* acquired DWV by feeding on adult bees which had become infected via food delivered to them by infected colony-mates. A group of 12 *Varroa* exposed to infectious (experimental) bees and 7 *Varroa* exposed to non-infectious (control) bees were individually sampled for presence of the tagged virus. Of the *Varroa* exposed to adult bees in the experimental group, 50% were positive for the novel virus (6/12 bees), while none of the controls reported detections (0/7 bees). Of the *Varroa* which reported detections, qPCR results showed low levels of the tagged virus (mean ± SD = 4.61log_10_ ± 0.17, n = 6). Natural DWV-A levels were not significantly different between exposed and unexposed *Varroa* in this study (AOV, *F*_*1,17*_ *=* 1.38, *p* = 0.26). 100% of *Varroa* in the experimental group had detections of DWV-A while 71.4% of *Varroa* which fed upon the unexposed bees had detections. (Table 2.1) After their removal from their adult bee hosts 9 of the 12 *Varroa* in the exposed group were transferred to feed upon pupae. (The other three *Varroa* were removed from their adult bee hosts and saved directly at -80°C until RNA extraction. These *Varroa* did not have a paired pupal host). Of the 9 *Varroa* with a paired pupal host, 5 of the *Varroa* had detectable levels of the Nluc RNA, while only one paired *Varroa*-pupal host also had detectable levels. These samples were screened for the tagged virus via the *Pac*1 restriction site and gel electrophoresis and detection was verified in both the *Varroa* and pupal host. 2 other positive detections of the NLuc RNA were observed in pupae not paired with *Varroa* samples. Overall observed transmission from *Varroa* to pupal hosts was 17.6% N = 17 (Supplementary Information). There was a general trend for less prevalence of the DWV pathogen after transmission cycles further away from the index host or index vector.

**Table 2.1.** Mean, median and SD of DWV-A and Nluc levels between experimentally exposed or control *Varroa*.

| DWV levels in pupae *Varroa* fed upon after feeding on adult bees (all values in log10 GE per bee) |  |  |  |  |  |  |
| --- | --- | --- | --- | --- | --- | --- |
| Trial | Group | State | Primer | Mean | Median | SD |
| Cana | Experimental | Naive | DWV | 6.34 | 6.25 | 2.08 |
| Cana | Control | Naive | DWV | 5.85 | 5.77 | 2.23 |
| Cana | NegControl | Naive | DWV | 5.50 | 4.83 | 2.06 |
| NLuc levels in pupae *Varroa* fed upon after feeding on adult bees (all values in log10 GE per bee) |  |  |  |  |  |  |
| Trial | Group | State | Primer | Mean | Median | SD |
| Cana | Experimental | Naive | NLuc | 4.60 | 5.86 | 2.60 |
| Cana | Control | Naive | NLuc | 0 | 0 | 0 |
| Cana | NegControl | Naive | NLuc | 0 | 0 | 0 |

### Experiment 2: Do Varroa acquire virus when feeding upon adult bees which previously cannibalized pupae?

Naïve *Varroa* transmitted the novel tagged virus to a pupal host after feeding upon adult bees which had cannibalized infectious pupae. Of 21 staged *Varroa*, 17 (80.9%) successfully transmitted the acquired tagged virus to a naïve bee host. No detections of the novel nLuc insert were observed in the control groups (n = 23). There was no significant difference in wild type DWV levels between experimentally exposed and control *Varroa* (AOV, *F*_*1,43*_ *=* 1.51, *p* = 0.29, Table 2.1)

### Experiments 3-4: Does *Varroa* to *Varroa* transmission occur on shared hosts?

Transmission occurred between *Varroa* that were concurrently feeding on pupal or adult bee hosts. Naïve *Varroa* acquired and subsequently transmitted a tagged virus while sharing a pupal host with another infectious *Varroa* (n = 20, 11/20 confirmed cases). There was no significant difference in wildtype or clone-derived virus loads between *Varroa* sharing a pupal host within the experimental group. (AOV, *F*_*1,37*_ *=* 0.042, *p* = 0.84 and *F*_*1,37*_ *=* 1.70, *p* = 0.2). Nor was there a significant difference in wildtype DWV-A levels between originally infectious and originally naïve *Varroa* within the experimental group which shared a pupal host (AOV, *F*_*1,37*_ *=*0.042, *p* = 0.84). DWV-A loads in these *Varroa* averaged 8.09 log_10_ and 9.93 log_10_ GE per bee in the two trials, while their non-*Varroa* exposed counterparts averaged 4.47 log_10_ and 3.54 log_10_ GE suggesting *Varroa* infestations were causing covert infections (AOV, *F*_*1,136*_ *=* 17.55, *p* < 0.0001). Of the 11 *Varroa*, 4 acquired tagged virus (<>7 log_10_ GE/*Varroa*). *Varroa* which shared adult bee hosts (Experiment 4) were also shown to acquire the tagged virus (66.7% n = 6). All *Varroa* that originally were exposed to the tagged virus through pupal feedings had detectable levels of the NLuc reporter sequence, with a modest (but non-significant, AOV, *F*_*1,10*_ *=* 14.373, *p* = 0.10) reduction in virus load for the naïve *Varroa* in mixed pairs. All *Varroa* in this trial had detectable levels of natural DWV-A (N = 18).

## Discussion

*Varroa*, the honey bee, and Deformed wing virus offer a complex vector-host-pathogen relationship where the pathogen moves among hosts via multiple routes^10,11^. Here we provide experimental evidence confirming several of these routes in honey bee colonies and showing their relative importance for the maintenance of DWV in both *Varroa* and bee hosts. These insights lead to a predictive and testable theoretical framework for assessing how interactive effects determine communicable and vectored transmission dynamics.

Most importantly, while multiple *Varroa* are rarely found on the same bee at the same time, they exhibit serial parasitism, shifting from host to host to feed^20^. Shared hosts offer rich opportunities for bidirectional material exchange between host and vector. Indeed, we show that naïve *Varroa* acquire DWV when they feed on either adult or pupal bees following the feeding by an infectious *Varroa*. This indirect *Varroa*-to-*Varroa* transmission confirms that *Varroa* are “hypodermic needles”, adeptly moving pathogens from host to host thanks to their promiscuous feeding. Vector-host exchanges of this sort are readily observed across arthropod vectors, most notably in ticks where co-feeding allows for non-systemic transmission between vectors^26,27^. To add another layer that is unique to social groups, *Varroa* also acquired DWV from non-parasitized bees that had themselves acquired this virus from nestmates via communicable transmission. Host to host transmission occurs through trophallaxis between adult bees, and our results strengthen and quantify previous observations that infectious hosts transmit DWV communicably to susceptible nestmates^12^. We were able to track viral transmission from introduction through multiple circulations between hosts and vectors. Our system of using tagged viruses shows that the original source of virus in workers was an infectious *Varroa* which fed upon an adult bee (Figure 2.1). This virus was then traced from parasitized bees to nestmates and ultimately to a new *Varroa* vector, fulfilling the parameters of Koch’s postulate^32^. While we show tagged viruses at each stage, we had anticipated greater levels of DWV amplification in bees which acquired the novel virus through trophallaxis (Supplementary Information). The experimental design could have impeded this result if the mesh screens in our design reduced trophallactic interactions. Alternatively, highly infectious bees might have died prior to phase two of the trial, impeding their ability to act as a maintenance host to susceptible nestmates. Such premature death of reservoir hosts is key for diminishing pathogen maintenance in a population and should be assessed in field trials^33^.

**Figure 2.1.**
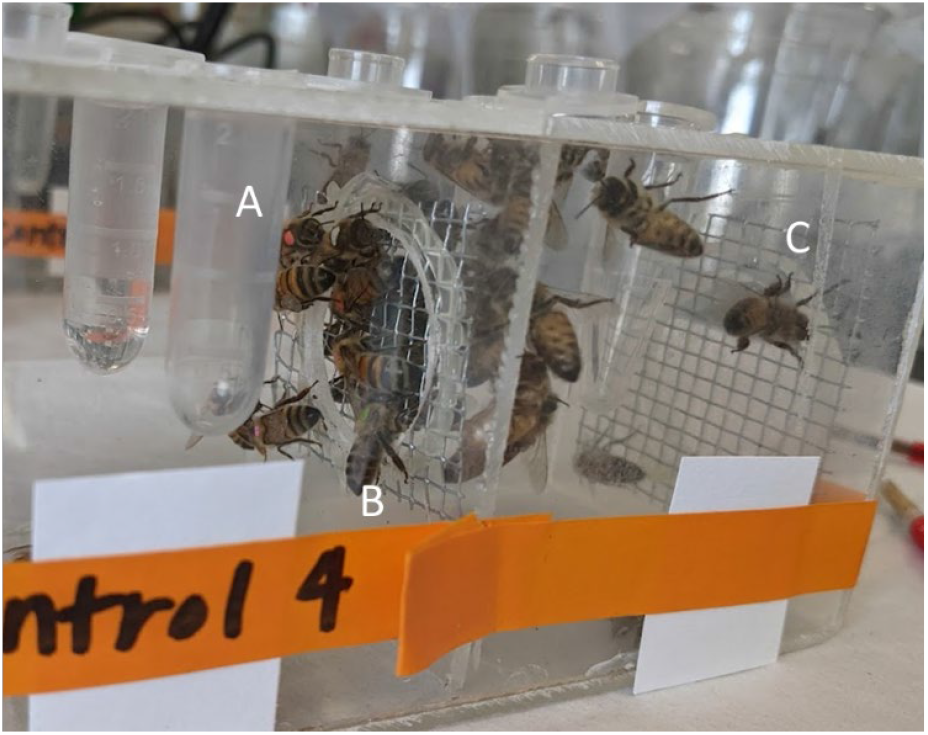
**(A)** Donor bee compartment. **(B)** Mesh screen divider. **(C)** Recipient bee compartment.

**Figure 2.2.**
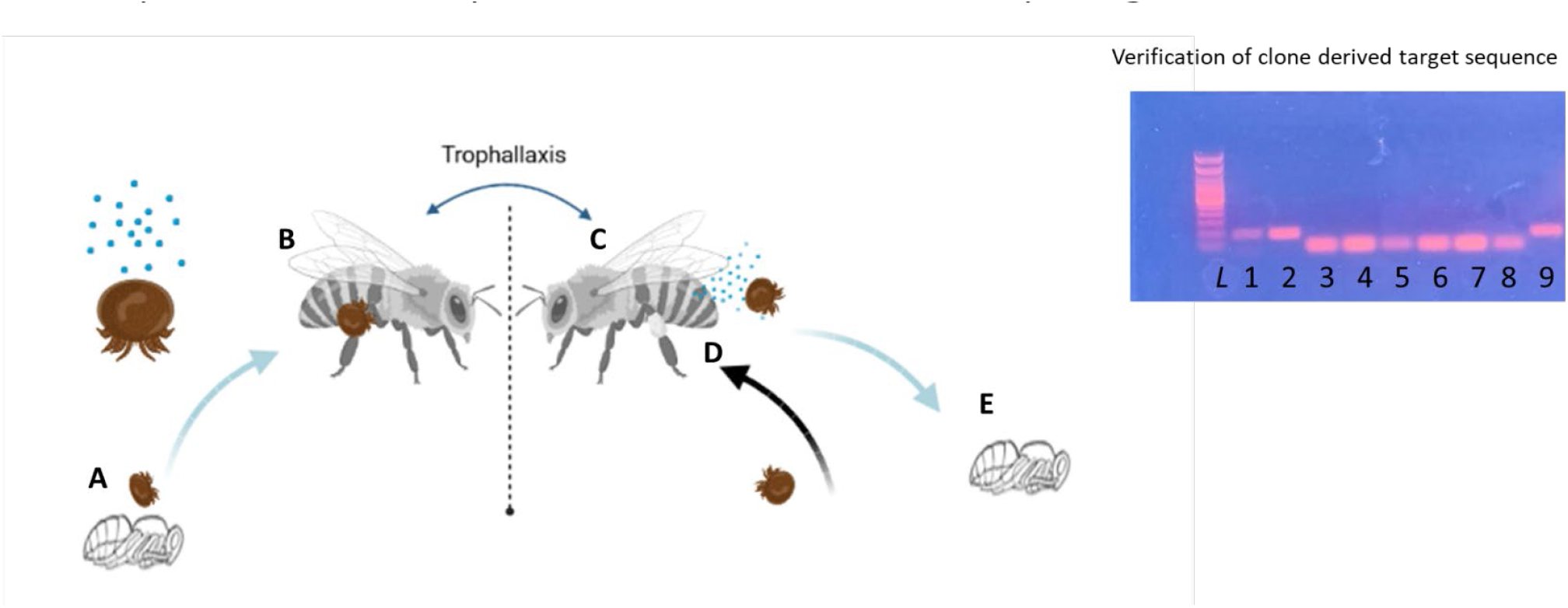
Transmission of DWV-A: **(A)** DWV-A (with NLuc reporter) was introduced into our system via infectious *Varroa*. (**B)** *Varroa* successfully vectored the pathogen to adult bee hosts who subsequently transmitted the pathogen through trophallaxis to naïve recipient bees **(C)** (unexposed to original *Varroa*). **(D)**New *Varroa* were introduced into the system which acquired the pathogen from the recipient bees, and **(E)** then vectored it to new pupal hosts. Detections of the tagged virus in *Varroa* mites was accomplished through the *Pac*I digestion of the 200 nt RT-PCR product containing the tagged region of DWV genome and visualization of the products by agarose gel electrophoresis. (**Right panel)** Lane L:100 nt DNA size ladder, **Lane 1** – *Varroa* “D” with both wildtype DWV (undigested) and the *Pac*I tagged DWV (faint digested bands are visible), Lane **2 -***Varroa* with wildtype DWV (no digestion), **Lanes 3,4** - paired samples: infectious *Varroa* (3) shared adult bee host with naïve *Varroa* (4); which both fed upon the same adult bee, Lane **5** - pupae (**E)** which was infected with the tagged virus by Varroa “D”. **Lanes 6,7 -** *Varroa* mites which shared the same pupal host (Lane 6 - was originally naïve Varroa lane 7 - infectious Varroa. Control honeybee infected with the tagged DWV-A: digested with PacI (line 8) and undigested (Line 9). Pupal image credit: D G Mackean www.biology-resources.com, Created with BioRender.com.

**Figure 2.3.**
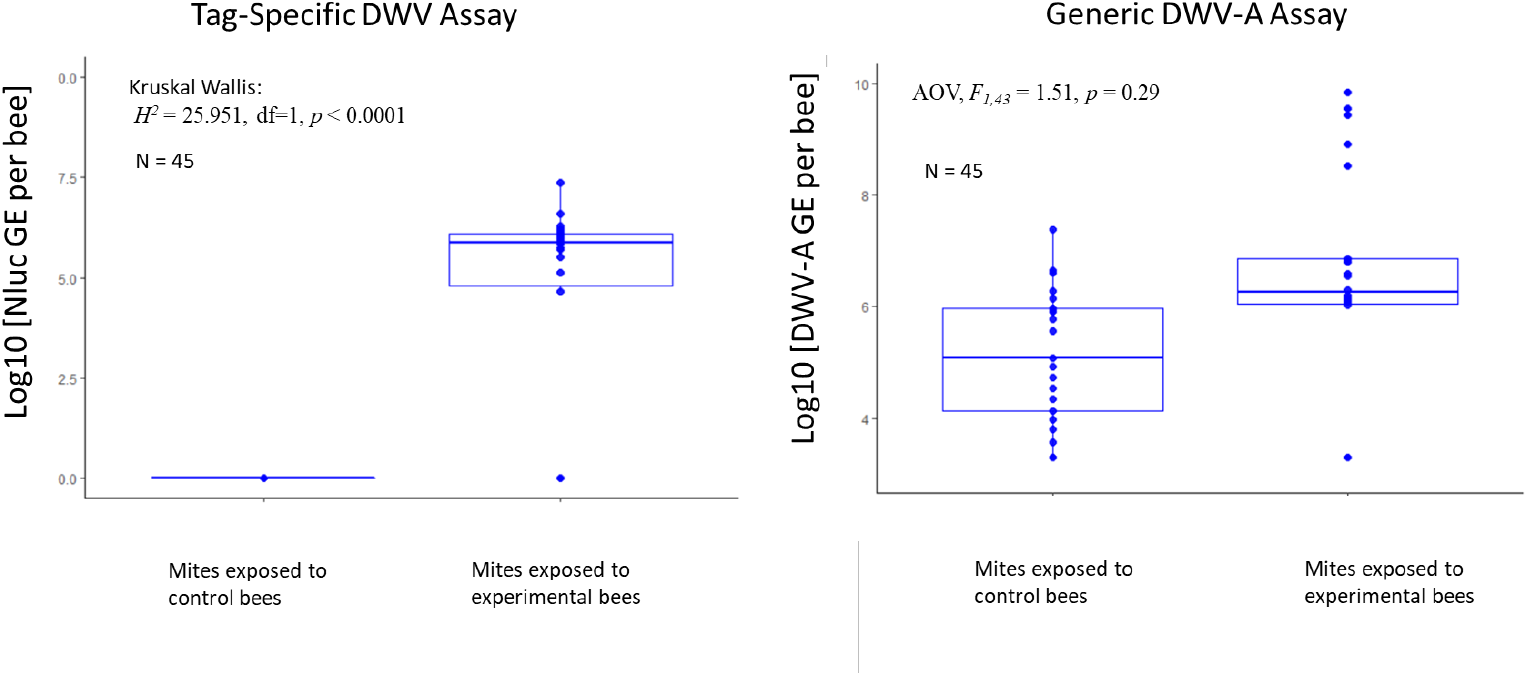
**(Left)** Detections of NLuc nucleotide sequence in pupae which mites fed upon after removal from their adult bee host. *Varroa* were exposed to either control or experimental adult bees. *Varroa* which fed on adult bees that cannibalized infectious pupae were highly likely to later transmit the tagged virus acquired through adult feeding (80.9%, N = 20). (**Right)** Detections of DWV-A in pupae which *Varroa* fed upon after removal from their adult bee host. There was no significant difference in natural DWV-A loads between control and experimental groups (AOV, *F*_*1,43*_ *=* 1.51, *p* = 0.29).

**Figure 2.4.**
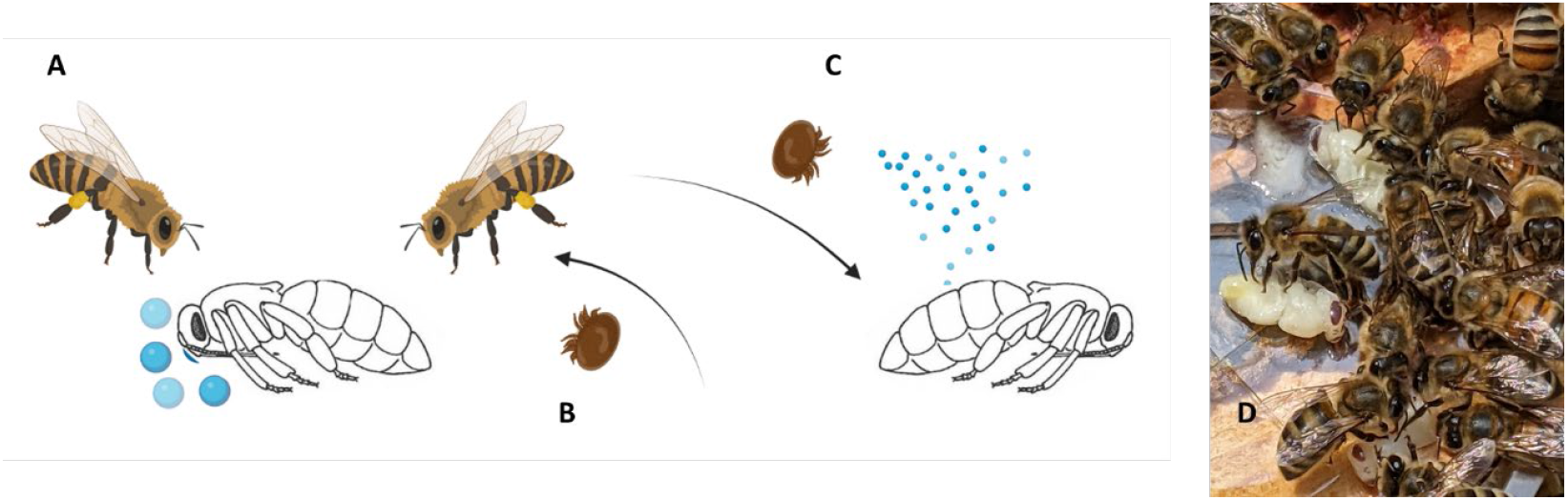
A schematic of transmission routes in experiment 2. **(A)** Adult worker bees cannibalize pupae infected with DWV-A NLuc. **(B)** *Varroa* mites feed on these adult bees. **(C)** Mites acquire and vector DWV NLuc into subsequent hosts. **(D)** Picture of adult worker bees inside of a colony cannibalizing pupae. (Pupal image credit: D G Mackean www.biology-resources.com, Created with BioRender.com.)

**Figure 2.5.**
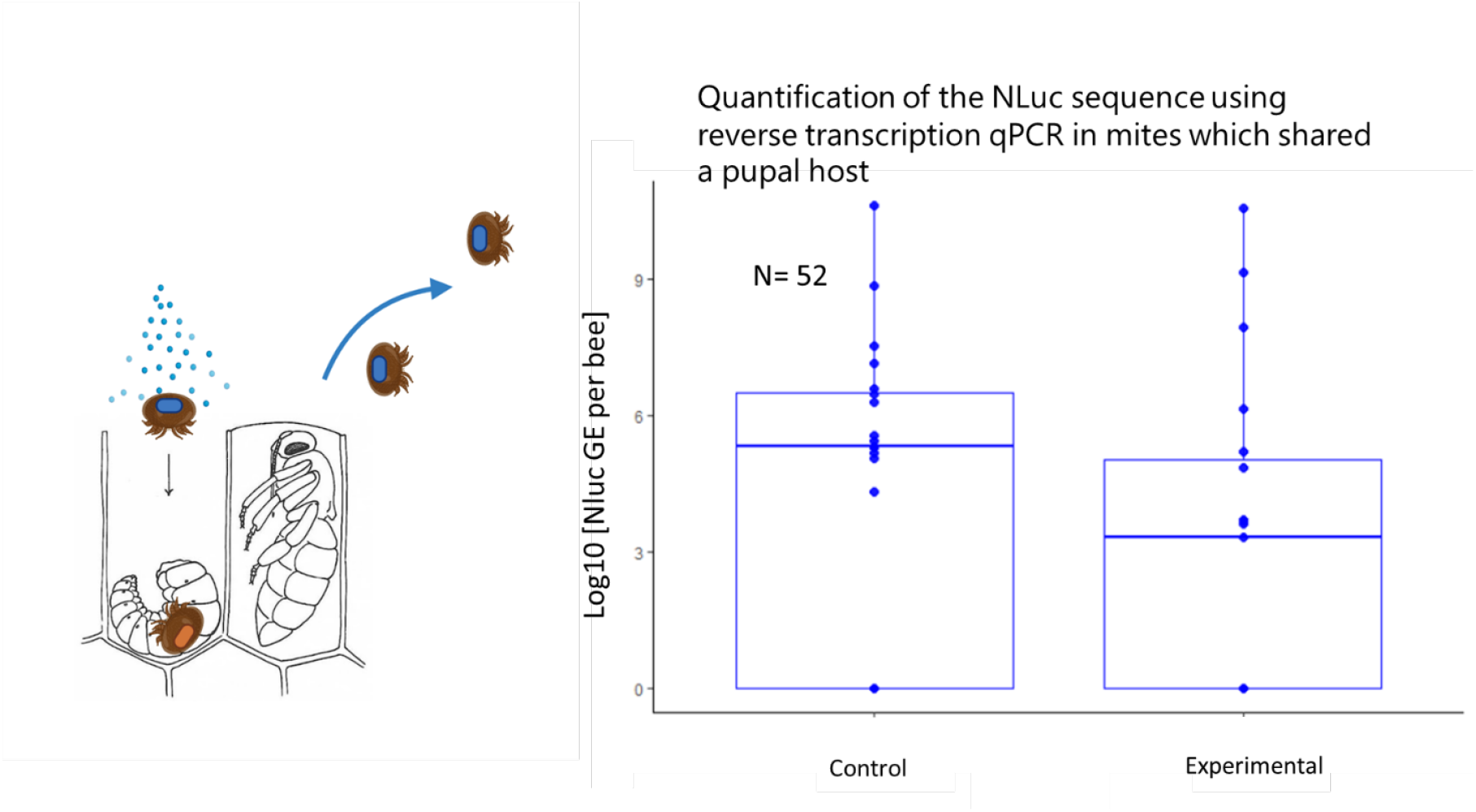
A schematic describing the conceptual framework of the study. Naïve *Varroa* may acquire infectious DWV-A when sharing the same pupal host with an infectious *Varroa. Varroa* which were originally infectious did not have significantly higher levels of DWV-NLuc compared to the originally naïve *Varroa* which shared a host. Fewer naïve *Varroa* transmitted the tagged virus than originally infectious *Varroa* which maintained the virus. (10/18 (55.56%) naïve successfully trasnmitted versus 14/21 (66.67%) combined from both trials. (Pupal image credit: D G Mackean www.biology-resources.com, Created with BioRender.com.)

In experiment two we asked if cannibalization, a social behavior common in honeybees as a route of removing infected nestmates or in times of nutritional stress, could provide a communicable route for transmission of DWV. We have previously shown that cannibalization of infectious pupae, coupled with oral transmission among nestmates through trophallaxis, could lead to viral transmission^12^. Here, we show that bees which cannibalize infectious pupae act as infectious hosts to naïve *Varroa* and that these *Varroa* subsequently vectored the novel DWV. These findings suggest that pupal cannibalization, which has been bred into honeybee colonies as a form of behavioral hygiene^12^, presents a possible risk factor. Such behavioral traits provide benefits by reducing *Varroa* reproduction and survival^34^, while likely increasing transmission of DWV to other nestmates. It is critical to assess, at the colony scale, the impacts of these opposing forces on colony health.

Although it was not within the scope of this study, there was an overall trend across experiments for the percentage of infectious vectors or hosts to be fewer after transmission cycles in the groups that did not originate with an index case. For example, at the end of experiment 1 there were more *Varroa* with detectable levels of DWV-A within the *Varroa* + virus group than either of the control groups (Supplementary Information). The *Varroa* + virus group began the experiment with an infectious vector, whereas the other groups did not. It is likely transmission efficiencies are not 100% effective. This would mean after subsequent *Varroa*-bee feedings or bee-bee trophallaxis cycles we would expect fewer infectious cases. Because we did not originally seek to answer this question our experimental design did not include a large enough number of observations to answer this question (Supplementary Information for count data on trials 1-4). The mechanisms behind this observation was outside the scope of this study, but warrants continued research in transmission efficiencies within this system.

DWV was present, but not prevalent, in honeybee populations before *Varroa destructor* jumped from *A. cerana* to *A. mellifera*^8,35,36^. Host-to-host transmission must have occurred to maintain DWV in honeybee populations prior to vectored transmission by *Varroa*. Host-to-host transmission has been attributed to vertical transmission of DWV from queen to progeny^14^. Our results indicate that worker-to-worker routes of transmission may rival those from queen to offspring. Since workers are targeted by *Varroa* more often than queens^37^, we would argue that worker-worker transmission is the predominant means of maintaining DWV in colonies and puts these colonies highly vulnerable when the vector arrives. Continued dispersal by these *Varroa* supports this, i.e., when *Varroa* arrive in an area, DWV levels in worker honey bees tend to increase exponentially and covary with *Varroa* infestation levels. When *Varroa* mites were introduced to the Hawaiian islands, viral loads in individual worker bees increased by a million-fold, establishing *Varroa*’s culpability in the emergence of DWV as a global pathogen^8^. The interplay of vector-host and host-host transmission for disease is rare in nature. In fact, the majority of infectious diseases are communicable, or vector borne, but very rarely both. Curiously, a select handful of emerging, RNA viruses, Zika, West Nile, Tembusu, salmon isavirus, Japanese encephalitis virus and DWV, are the exceptions to this rule^4,10,38-40^. As our results show, understanding this interplay can offer fundamental insights into routes of transmission and disease, and can point to management strategies that might break transmission, improving the health of a key pollinator.

## Author Contributions

ZSL conceptualized the project and methodology. ZSL performed the investigation and statistical analysis. EVR supervised and oversaw viral work and analysis. ZSL, JDE, DJH: original draft preparation. ZSL, EVR, DJH, JDE: writing-review and editing. DJH and JDE: supervision.

## References

1 Grassly, N. C. & Fraser, C. Mathematical models of infectious disease transmission. Nature Reviews Microbiology 6, 477–487, doi:10.1038/nrmicro1845 (2008).

2 Anderson, R. & May, R. Infection diseases of humans. Dynamics and control. Oxford and New York: Oxford University Press (1991).

3 Garrett-Jones, C. THE HUMAN BLOOD INDEX OF MALARIA VECTORS IN RELATION TO EPIDEMIOLOGICAL ASSESSMENT. Bull World Health Organ 30, 241–261 (1964).

4 Hartemink, N. A., Davis, S. A., Reiter, P., Hubálek, Z. & Heesterbeek, J. A. Importance of bird-to-bird transmission for the establishment of West Nile virus. Vector Borne Zoonotic Dis 7, 575–584, doi:10.1089/vbz.2006.0613 (2007).

5 van den Driessche, P. Reproduction numbers of infectious disease models. Infectious Disease Modelling 2, 288–303, doi:https://doi.org/10.1016/j.idm.2017.06.002 (2017).

6 Wilfert, L. et al Deformed wing virus is a recent global epidemic in honeybees driven by <em>Varroa</em> mites. Science 351, 594–597, doi:10.1126/science.aac9976 (2016).

7 Martin, S. J. & Brettell, L. E. Deformed Wing Virus in Honeybees and Other Insects. Annual Review of Virology 6, 49–69, doi:10.1146/annurev-virology-092818-015700 (2019).

8 Martin, S. J. et al Global Honey Bee Viral Landscape Altered by a Parasitic Mite. Science 336, 1304–1306, doi:10.1126/science.1220941 (2012).

9 Oldroyd, B. P. What’s Killing American Honey Bees? PLoS Biology 5, e168, doi:10.1371/journal.pbio.0050168 (2007).

10 Chen, Y., Evans, J. & Feldlaufer, M. Horizontal and vertical transmission of viruses in the honey bee, Apis mellifera. Journal of Invertebrate Pathology 92, 152–159, doi:https://doi.org/10.1016/j.jip.2006.03.010 (2006).

11 Möckel, N., Gisder, S. & Genersch, E. Horizontal transmission of deformed wing virus: pathological consequences in adult bees (Apis mellifera) depend on the transmission route. Journal of General Virology 92, 370–377, doi:https://doi.org/10.1099/vir.0.025940-0 (2011).

12 Posada-Florez, F. et al Pupal cannibalism by worker honey bees contributes to the spread of deformed wing virus. Scientific Reports 11, 8989, doi:10.1038/s41598-021-88649-y (2021).

13 Yue, C. & Genersch, E. RT-PCR analysis of Deformed wing virus in honeybees (Apis mellifera) and mites (Varroa destructor). Journal of General Virology 86, 3419–3424, doi:https://doi.org/10.1099/vir.0.81401-0 (2005).

14 Amiri, E. et al Quantitative patterns of vertical transmission of deformed wing virus in honey bees. PLOS ONE 13, e0195283, doi:10.1371/journal.pone.0195283 (2018).

15 Amiri, E., Meixner, M. D. & Kryger, P. Deformed wing virus can be transmitted during natural mating in honey bees and infect the queens. Scientific Reports 6, 33065, doi:10.1038/srep33065 (2016).

16 Yue, C., Schroder, M., Gisder, S. & Genersch, E. Vertical-transmission routes for deformed wing virus of honeybees (Apis mellifera). Journal of General Virology 88, 2329–2336, doi:10.1099/vir.0.83101-0 (2007).

17 De Miranda, J. R. & Fries, I. Venereal and vertical transmission of deformed wing virus in honeybees (Apis mellifera L.). Journal of Invertebrate Pathology 98, 184–189, doi:10.1016/j.jip.2008.02.004 (2008).

18 Kanbar, G. & Engels, W. Ultrastructure and bacterial infection of wounds in honey bee (Apis mellifera) pupae punctured by Varroa mites. Parasitology Research 90, 349–354, doi:10.1007/s00436-003-0827-4 (2003).

19 Ramsey, S. D. et al Varroa destructor feeds primarily on honey bee fat body tissue and not hemolymph. Proceedings of the National Academy of Sciences 116, 1792–1801, doi:10.1073/pnas.1818371116 (2019).

20 Lamas, Z. S. Feeding behavior and dristribution of Varroa destructor on adult bees of Apis mellifera (University of Maryland, 2022).

21 Gisder, S., Aumeier, P. & Genersch, E. Deformed wing virus: replication and viral load in mites (Varroa destructor). Journal of General Virology 90, 463–467, doi:10.1099/vir.0.005579-0 (2009).

22 Yang, X. & Cox-Foster, D. L. Impact of an ectoparasite on the immunity and pathology of an invertebrate: Evidence for host immunosuppression and viral amplification. Proceedings of the National Academy of Sciences 102, 7470–7475, doi:10.1073/pnas.0501860102 (2005).

23 Sumpter, D. J. T. & Martin, S. J. The dynamics of virus epidemics inVarroa-infested honey bee colonies. Journal of Animal Ecology 73, 51–63, doi:10.1111/j.1365-2656.2004.00776.x (2004).

24 Bowen-Walker, P. L., Martin, S. J. & Gunn, A. The Transmission of Deformed Wing Virus between Honeybees (Apis melliferaL.) by the Ectoparasitic MiteVarroa jacobsoniOud. Journal of Invertebrate Pathology 73, 101–106, doi:10.1006/jipa.1998.4807 (1999).

25 Nordström, S. Distribution of deformed wing virus within honey bee (Apis mellifera) brood cells infested with the ectoparasitic mite Varroa destructor. Experimental & Applied Acarology 29, 293–302, doi:10.1023/A:1025853731214 (2003).

26 Labuda, M. et al Tick-Borne Encephalitis Virus Transmission between Ticks Cofeeding on Specific Immune Natural Rodent Hosts. Virology 235, 138–143, doi:10.1006/viro.1997.8622 (1997).

27 Labuda, M., Jones, L. D., Williams, T., Danielova, V. & Nuttall, P. A. Efficient transmission of tick-borne encephalitis virus between cofeeding ticks. Journal of medical entomology 30, 295–299 (1993).

28 Ogden, N., Nuttall, P. & Randolph, S. Natural Lyme disease cycles maintained via sheep by co-feeding ticks. Parasitology 115, 591–599 (1997).

29 Evans, J. D., Banmeke, O., Palmer-Young, E. C., Chen, Y. & Ryabov, E. V. Beeporter: Tools for high-throughput analyses of pollinator-virus infections. Mol Ecol Resour, doi:10.1111/1755-0998.13526 (2021).

30 Evans, J. D. et al Standard methods for molecular research inApis mellifera. Journal of Apicultural Research 52, 1–54, doi:10.3896/ibra.1.52.4.11 (2013).

31 Posada-Florez, F. et al Deformed wing virus type A, a major honey bee pathogen, is vectored by the mite Varroa destructor in a non-propagative manner. Scientific Reports 9, doi:10.1038/s41598-019-47447-3 (2019).

32 Rivers, T. M. Viruses and Koch’s postulates. Journal of bacteriology 33, 1 (1937).

33 Roberts, M. G. & Heesterbeek, J. A. P. Characterizing reservoirs of infection and the maintenance of pathogens in ecosystems. Journal of The Royal Society Interface 17, 20190540, doi:doi:10.1098/rsif.2019.0540 (2020).

34 Harris, J. W. Bees with Varroa Sensitive Hygiene preferentially remove mite infested pupae aged≤ five days post capping. Journal of Apicultural Research 46, 134–139 (2007).

35 De Souza, F. S., Allsopp, M. H. & Martin, S. J. Deformed wing virus prevalence and load in honeybees in South Africa. Archives of Virology 166, 237–241, doi:10.1007/s00705-020-04863-5 (2021).

36 Roberts, J. M. K., Anderson, D. L. & Durr, P. A. Absence of deformed wing virus and Varroa destructor in Australia provides unique perspectives on honeybee viral landscapes and colony losses. Scientific Reports 7, doi:10.1038/s41598-017-07290-w (2017).

37 Santillán-Galicia, M. T., Otero-Colina, G., Romero-Vera, C. & Cibrián-Tovar, J. Varroa destructor (Acari: Varroidae) infestation in queen, worker, and drone brood of Apis mellifera (Hymenoptera: Apidae). The Canadian Entomologist 134, 381–390, doi:10.4039/Ent134381-3 (2002).

38 Fish, I.-I. INF ECTIOUSNESS OF ORGANIC MATERIALS ORIGINAT-ING IN ISA-INFECTED FISH AND TRANSMISSION OF THE DISEASE VIA SALMON LICE (LEPE OPH THEIR US SAL-MONIS). Bull. Eur. Ass. Fish Pathol 18, 173 (1998).

39 Magalhaes, T., Foy, B. D., Marques, E. T. A., Ebel, G. D. & Weger-Lucarelli, J. Mosquito-borne and sexual transmission of Zika virus: Recent developments and future directions. Virus Research 254, 1–9, doi:10.1016/j.virusres.2017.07.011 (2018).

40 Cao, Z. et al Tembusu Virus in Ducks, China. Emerging Infectious Diseases 17, 1873–1875, doi:10.3201/eid1710.101890 (2011).

