## Supplemental Material for "Oversharing by honey bees and the spread of viruses"

#### Supplementary Information

A. Count data for Experiments 1,3,4.

#### Experiment 1

##### Mean and standard deviation of recipient bees per group that died during the trial

| Group | Total Recipients Added | Mortality | Average(STDEV) | AOV Mortality P (f) |
|---|---|---|---|---|
| *Varroa* + Virus | 150 | 13 | 2.6 (2.70) | P = .015 (2.223) |
| *Varroa* | 150 | 15 | 3 (2.92) | - |
| No *Varroa* | 150 | 30 | 6 (2.74) | - |

**Number of *Varroa* individually sampled for presence of NanoLuc**

| Group | Total # *Varroa* | Number detected | Number not detected | Percent of samples w/ detection |
|---|---|---|---|---|
| *Varroa* + Virus | 12 | 6 | 6 | 50% |
| *Varroa* | 3 | 0 | 3 | 0% |
| No *Varroa* | 4 | 0 | 4 | 0% |

**Number of *Varroa* individually sampled for presence of DWV-A**

| Group | Total # *Varroa* | Number detected | Number not detected | Percent of samples w/ detection |
|---|---|---|---|---|
| *Varroa* + Virus | 12 | 12 | 12 | 100% |
| *Varroa* | 3 | 2 | 1 | 67% |
| No *Varroa* | 4 | 3 | 1 | 75% |

**Number of pupae individually sampled for presence of NanoLuc**

*These are pupae a Varroa fed upon in the final stage of the experiment*

| Group | Total # *Varroa* | Number detected | Number not detected | Percent of samples w/ detection |
|---|---|---|---|---|
| *Varroa* + Virus | 17 | 3 | 14 | 17.6% |
| *Varroa* | 14 | 0 | 14 | 0% |
| No *Varroa* | 6 | 0 | 6 | 0% |

**Number of pupae individually sampled for presence of DWV-A**

*These are pupae a Varroa fed upon in the final stage of the experiment*

| Group | Total # *Varroa* | Number detected | Number not detected | Percent of samples w/ detection |
|---|---|---|---|---|
| *Varroa* + Virus | 17 | 16 | 1 | 94.1% |
| *Varroa* | 14 | 11 | 3 | 78.6% |
| No *Varroa* | 6 | 4 | 2 | 66.7% |

#### Experiment 1 (continued)

**Mean and standard deviation of donor bees per treatment (replicates, n = 5)**

| Treatment | Total Donors Added | Mortality | Average(STDEV) | AOV Mortality P(F) |
|---|---|---|---|---|
| *Varroa* + Virus | 74 | 20 | 4 (1) | P = .08 (3.206) |
| *Varroa* | 64 | 15 | 3 (1.87) | - |
| No *Varroa* | 75 | 8 | 1.8 (1.41) | - |

**Count data for viral detections in naïve *Varroa* which fed on recipient bees**

| Group | Target | Sample Type | Total Samples | Number detected | Number not detected | Percent of samples w/ detection |
|---|---|---|---|---|---|---|
| *Varroa* + Virus | NanoLuc | *Varroa* | 12 | 6 | 6 | 50% |
| *Varroa* | NanoLuc | *Varroa* | 3 | 0 | 3 | 0% |
| No *Varroa* | NanoLuc | *Varroa* | 4 | 0 | 4 | 0% |
| Experimental | DWV-A | *Varroa* | 12 | 12 | 12 | 100% |
| *Varroa* + Virus | DWV-A | *Varroa* | 3 | 1 | 2 | 67% |
| *Varroa* | DWV-A | *Varroa* | 4 | 3 | 1 | 75% |
| *Varroa* + Virus | NanoLuc | Pupa | 17 | 3 | 14 | 17.6% |
| *Varroa* | NanoLuc | Pupa | 14 | 0 | 14 | 0% |
| No *Varroa* | NanoLuc | Pupa | 6 | 0 | 6 | 0% |
| Experimental | DWV-A | Pupa | 17 | 16 | 1 | 94.1% |
| *Varroa* + Virus | DWV-A | Pupa | 14 | 11 | 3 | 78.6% |
| *Varroa* | DWV-A | Pupa | 6 | 4 | 2 | 66.7% |

#### Experiment 3: Co-Feeding on brood

**In This trial, Cofeeding on brood, CF (June, 2022) 19 hosts housed 38 *Varroa*. *Varroa* were divided into two groups (experimental (n = 25) and control (n = 6)**

**Recollection of *Varroa* and detection of molecular targets in pupae they fed upon after recollection**

| Group | Target | Total # recollected *Varroa* | Number pupae detected with NanoLuc | Number pupae without detection | Percent of samples w/ detection |
|---|---|---|---|---|---|
| Experimental-Naïve | NanoLuc | 11 | 6 | 5 | 54.5%% |
| Experimental Infectious | NanoLuc | 14 | 8 | 6 | 57.1% |
| Control | NanoLuc | 6 | 0 | 6 | 0% |
| Experimental-Naïve | DWV | 11 | 10 | 1 | 90.9%% |
| Experimental Infectious | DWV | 14 | 13 | 1 | 92.9% |
| Control | DWV | 6 | 5 | 1 | 83.3% |

**Recollection of *Varroa* and detection of DWV-A in pupae they fed upon after recollection**

| Group | Total # recollected *Varroa* | Number pupae detected with DWV-A | Number pupae without detection | Percent of samples w/ detection |
|---|---|---|---|---|
| Experimental-Naïve | 11 | 10 | 1 | 90.9%% |
| Experimental Infectious | 14 | 13 | 1 | 92.9% |
| Control | 6 | 5 | 1 | 83.3% |

#### Experiment 3: Co-Feeding on brood (continued)

**Viral loads in CF-(Cofeeding-Brood Trial, June 2022)**

| Trial | Group | State | Primer | Mean | Median | SD |
|---|---|---|---|---|---|---|
| CF | Experimental | All Samples | DWV | 8.090696 | 7.951268 | 2.393663 |
| CF | Control | All Samples | DWV | 6.125059 | 5.278225 | 2.750206 |
| CF | Experimental | Infectious | DWV | 8.004581 | 7.891846 | 2.36842 |
| CF | Experimental | Naive | DWV | 8.200296 | 7.951268 | 2.536781 |
| CF | Control | All | LUC | No detection | No detection | Nodetection |
| CF | Experimental | All | LUC | 3.783416 | 3.707195 | 3.811959 |
| CF | Experimental | Infectious | LUC | 4.314553 | 5.350022 | 3.836407 |
| CF | Experimental | Naive | LUC | 3.208017 | 1.845109 | 3.866971 |
| CF | Background Control Pupae: Start | | DWV | 5.869487 | 5.896935 | 0.3925235 |
| CF | Background Control Pupae: Middle | | DWV | 4.455652 | 4.295666 | 1.283693 |
| CF | Background Control Pupae: End | | DWV | 4.471733 | 4.424042 | 1.284144 |

**These include only samples with a positive detection of LUC and excludes all zeros (No detects)**

| Trial | Group | State | Primer | Mean | Median | SD |
|---|---|---|---|---|---|---|
| CF | Experimental | Infectious | LUC | 7.011149 | 6.396124 | 1.903618 |
| CF | Experimental | Naive | LUC | 6.416034 | 5.689485 | 2.863271 |

**Trial: Co-feeding on brood (CFA, August, 2021)**

**In This trial, Cofeeding on brood, CF (July, 2022) 24 hosts housed 48 *Varroa*. *Varroa* were divided into two groups (experimental (n = 24) and control (n = 24).**

**Recollection of *Varroa* and detection of NLuc in pupae they fed upon after recollection**

| Group | Total # recollected *Varroa* | Number pupae detected with NanoLuc | Number pupae without detection | Percent of samples w/ detection |
|---|---|---|---|---|
| Experimental- Naïve | 7 | 4 | 3 | 57.1%% |
| Experimental Infectious | 7 | 6 | 1 | 85.7% |
| Control | 14 | 0 | 6 | 0% |

**Recollection of *Varroa* and detection of DWV-A in pupae they fed upon after recollection**

| Group | Total # recollected *Varroa* | Number pupae detected with DWV-A | Number pupae without detection | Percent of samples w/ detection |
|---|---|---|---|---|
| Experimental- Naïve | 7 | 7 | 0 | 100% |
| Experimental Infectious | 7 | 7 | 0 | 100% |
| Control | 14 | 0 | 0 | 100% |

#### Experiment 3: Co-Feeding on brood (continued)

**Viral loads in CFA-(Cofeeding-Brood Trial, August 2022)**

| Trial | Group | State | Primer | Mean | Median | SD |
|---|---|---|---|---|---|---|
| CFA | Experimental | All Samples | DWV | 9.926413 | 10.7184 | 1.436755 |
| CFA | Control | All Samples | DWV | 9.180214 | 9.169762 | 1.566407 |
| CFA | Experimental | Infectious | DWV | 10.03198 | 10.74724 | 1.336651 |
| CFA | Experimental | Naive | DWV | 9.820845 | 10.30884 | 1.630931 |
| CFA | Experimental | All Samples | LUC | 3.827413 | 4.596895 | 2.799905 |
| CFA | Control | All Samples | LUC | No detection | No detection | Nodetection |
| CFA | Experimental | Infectious | LUC | 4.827161 | 5.17646 | 2.334959 |
| CFA | Experimental | Naive | LUC | 2.827665 | 3.345055 | 3.033367 |
| CFA | Background Control Pupae: Start | | DWV | 4.247459 | 4.243642 | 0.3959113 |
| CFA | Background Control Pupae: Middle | | DWV | 4.236862 | 4.183093 | 0.2607861 |
| CFA | Background Control Pupae: End | | DWV | 3.538158 | 3.547232 | 0.528503 |

**These include only samples with a positive detection of LUC and excludes all zeros (No detects)**

| Trial | Group | State | Primer | Mean | Median | SD |
|---|---|---|---|---|---|---|
| CFA | Experimental | Infectious | LUC | 5.631688 | 5.310558 | 1.051393 |
| CFA | Experimental | Naive | LUC | 4.948414 | 4.25416 | 2.100042 |

(continued on next page)

### Experiment 4: Co-feeding on adult bees

**The number of *Varroa* collected and individually extracted for NLuc**

| Group | Total # *Varroa* | Number detected | Number not detected | Percent of samples w/ detection |
|---|---|---|---|---|
| Experimental | 12 | 12 | 12 | 100% |
| Control | 6 | 5 | 2 | 41.7% |
| Experimental Naive | 6 | 4 | 2 | 66.7% |
| Experimental Infectious | 6 | 6 | 0 | 100% |

**The number of *Varroa* collected and individually extracted for DWV-A**

| Group | Total # *Varroa* | Number detected | Number not detected | Percent of samples w/ detection |
|---|---|---|---|---|
| Experimental | 12 | 12 | 12 | 100% |
| Control | 6 | 6 | 0 | 100% |
| Experimental Naive | 6 | 4 | 2 | 100% |
| Experimental Infectious | 6 | 6 | 0 | 100% |
